# Data Reuse and the Social Capital of Open Science

**DOI:** 10.1101/093518

**Authors:** Bradly Alicea

**Affiliations:** Orthogonal Research and Education Laboratory, Champaign IL, USA http://orthogonal-research.weebly.com

**Keywords:** Game Theory, Open Science, Data Reuse, Econosemantics

## Abstract

Participation in open data initiatives require two semi-independent actions: the sharing of data produced by a researcher or group, and a consumer of shared data. Consumers of shared data range from people interested in validating the results of a given study to people who actively transform the available data. These data transformers are of particular interest because they add value to the shared data set through the discovery of new relationships and information which can in turn be shared with the same community. The complex and often reciprocal relationship between producers and consumers can be better understood using game theory, namely by using three variations of the Prisoners’ Dilemma (PD): a classical PD payoff matrix, a simulation of the PD *n*-person iterative model that tests three hypotheses, and an Ideological Game Theory (IGT) model used to formulate how sharing strategies might be implemented in a specific institutional culture. To motivate these analyses, data sharing is presented as a trade-off between economic and social payoffs. This is demonstrated as a series of payoff matrices describing situations ranging from ubiquitous acceptance of Open Science principles to a community standard of complete non-cooperation. Further context is provided through the IGT model, which allows from the modeling of cultural biases and beliefs that influence open science decision-making. A vision for building a CC-BY economy are then discussed using an approach called econosemantics, which complements the treatment of data sharing as a complex system of transactions enabled by social capital.

## Introduction

In a rather short period of time, the culture and practice of Open Science [1] has contributed to substantial changes in academic practices. One way to assess the advantages of openly sharing data sets, both in terms of scientific progress and diffusion of scientific knowledge is to use a formal model. It is proposed here that sharing and consuming open data can be modeled as a Prisoners’ Dilemma (PD) game. In a basic PD game, two players can either cooperate or defect (not cooperate) given information about their partner’s decision. There strategies can be employed in a pure fashion (one player always cooperate or defect) or in a mixed fashion where players alternate their strategy over time. As an analogy to data sharing, cooperation entails a producer of data sharing their data with another member of their community. The community members transform (actively consumer) the data into publications, meta-analyses, new knowledge, or new data sets. By contrast, defection is analogous to the withholding of data by producers, or a failure to reciprocate by the data transformers. Reciprocation is broader than simply producing data to share with the community. It can include citation ot co-authorship as a means to strengthen the incentive for future sharing.

The classical PD game, PD *n*-player iterative model, and PD modeled using a hybrid game-theoretic/mathematical model called Ideological Game Theory (IGT) lend new insights into the role of cooperation [2], non-cooperation, and altruism [3] in open data communities. In addition to empirical investigations of these communities, researchers have also begun to view “open” scholarly transactions as a problem of joint cooperation and an appropriate application of mathematical and game theory models. Pronk et.al [4] introduced a mathematical model for the cost in time and resources of optimal reuse conditions with regard to impact. The associated argument for the model’s validity focuses on the legal, financial, and strategic costs, while this paper focuses on the hypothetical sociocultural conditions conducive to reuse [5]. A few other applications of PD models to open science and open access more broadly make some predictions about the nature of joint cooperation. For example, the PD model has been applied to the adoption of open access publications [6] to demonstrate “slow scalability” (the rate of adoption and reach within a scholarly community). Scheliga and Friesike [7] use a PD model to identify two obstacles to sustainable open communities: individual and systematic. Additionally, the application of a PD model provides a means for understanding diversity in the data sharing community [8]. In particular, when heterogeneous strategies are employed by community member (cooperate, defect, a mix of cooperate and defect, or some other conditional strategy), the outcome across the social network favor neither cooperation nor defection. Overall, the practice of openness and its associated benefits must be consistent across both the producers of primary data and the transformers of secondary data (Figure 1). While these acts may be independent of each other, this type of system must involve shared objectives to be sustainable.

**Figure 1.**
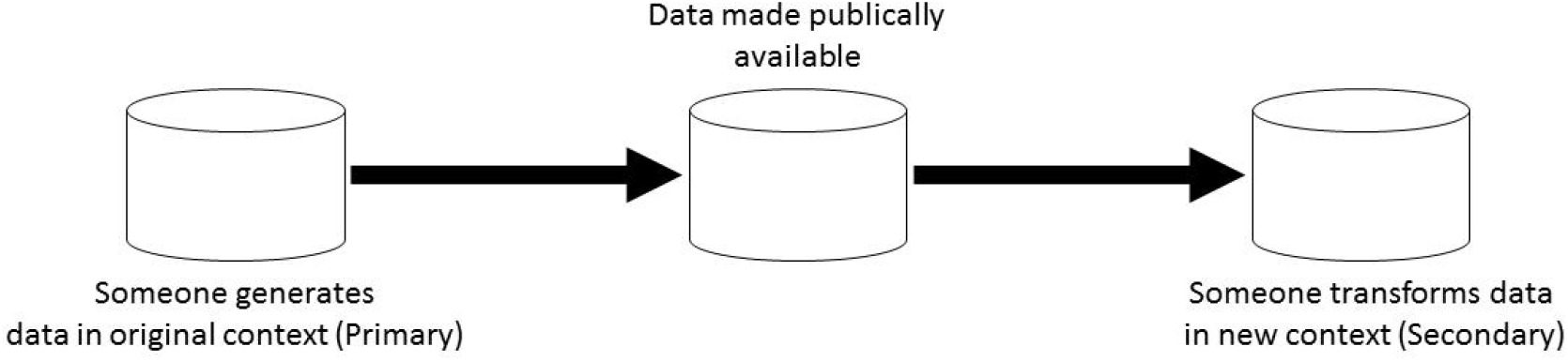
Typical pipeline of shared data between primary generation and secondary transformation.

### Prior Work and Background

To formulate this set of interactions in a series of theoretical models, we must determine the payoff structure of resulting from the relative openness and associated practices in a community of data sharers. We can justify our selection of theoretical payoff structures in the current study by looking to both previous game-theoretic and empirical studies on the topic. One way to approximate these practices is to incorporate surveys of both primary data producers and secondary data consumers. Taken collectively, this information to determine their existing habits and cultural mores. A survey of author behavior in light of enforcing sharing norms on producers has been examined using *Nature* journal submissions by [9]. In this case, imposing sharing standards on producers results in a time investment on the part of the producer, which in turn may constrain the extent of sharing on the part of said producer. Another indicator of how habits, mores, and cultural constraints influence payoffs is to examine cases in which producers can choose their level of contribution. An existing game-theoretic model [10] suggests that the more trustworthy and precise the data generated by producers, the more likely an equilibrium will develop between producers and consumers. When participants in a user-generated content (UGC) community can choose their degree of contribution, they tend to be highly motivated in their subsequent participation [11]. As with the case of UGCs, there are also parallels between an emerging data commons and the creation of public goods [12]. Viewed as a public good, the formation of a data commons relies heavily on producers controlling the payoff structure for the entire community, but in a way that ensures both shared costs and mutual payoffs for both producers and consumers.

Empirical studies of ad-hoc data sharing [13] can also help us understand under what conditions selfish and altruistic behaviors are exhibited. These stand in contrast to cooperative behaviors in that they are not particularly bound by social conventions. This is studied in a simulation of 200 users with a focus on non-selfish behaviors. For a strategy of reciprocation, the proportion of selfish users increases when the number of selfish users is less than 50. In cases where an altruistic strategy is employed, there is a net benefit for altruists even when a relatively low number of non-selfish users exist in the community. Is this a matter of shared cultural mores, or can openness be achieved merely through community enforcement of standard practices through incentives? To understand this better, it can be proposed that the enabling factor for openness is based on social rather than financial transactions. Social transactions are quantifiable using the concept of social capital [14]. Favorable social interactions result in increased social capital, while discouraged social interactions result in a loss of social capital [15]. Conforming to the prevailing social norms, or producing something that helps others within that set of norms, can produce social capital-derived utility for the transactor. This is distinct from financial capital, but operates in conjunction with social capital in order to determine the utility of engaging in data sharing and reuse.

### Outline of Approach

This paper is organized in the following manner. First, we will define some assumptions of the game-theoretic approach. These include making connections between topics in open data and how they are empirically realized in the academic context. The methods for our approach are presented next, which include the technical details of an agent-based model in addition to the experimental design, computational hypothesis, and payoffs of the PD approach. Next is a theoretical discussion of applying the PD game to data sharing, with particular focus on how behaviors and equilibria typically associated with PD is realized in an academic context. Then we report the results of an agent-based model of the PD game (the PD *n*-player iterative model) and discuss the results of these investigations. The simulation is followed up with a more speculative theoretical formulation that proposes using IGT as a means to better understand the epistemological variation that affects cultural practice and change within specific fields. In conclusion, the limitations of our approach and current state of knowledge are presented, as well as a consideration of how this work might contribute to the development of an economic model (econosemantics) organized around Creative Commons principles.

### Assumptions

Before we can calculate our payoffs, we must clarify a few assumptions. The first assumption is one of openness: what is the incentive for a producer to share data? While there are an increasing number of institutional encouragements for making access to research materials open [16], there still exists heterogeneity of practice in light of top-down incentives such as data availability statements [9]. There should instead be a focus on the relationship between time, financial gain, and utility. Sharing data requires an investment of time in data organization, annotation, and the presence of an easy-to-use repository that allows for the assignation of credit [17, 18]. One concrete example of this is an increase in citations for publications that make their data open [19]. In particular, professional credit can be thought of as social capital, and plays a role in the utility of sharing data [20]. This includes (but is not limited to): recognition by tenure and promotion committees, normative recognition of Altmetric milestones [21], and an increased recognition of reanalysis and replication studies as a valid form of scientific discovery [22].

One way to remove the effects of immediate financial incentive is to assume that all data are completely open access. In this case, completely open access means free of subscription and publication fees, or Gold Open Access without an embargo period as per [1]. To assume that open data is superior to non-open or withheld data, the following relation must hold: the utility (social capital) gained from sharing data must exceed the utility gained from withholding data or limiting access to one’s immediate collaborators. While this seems to be a difficult calculation to make, sharing data has a number of obvious benefits. Not only does open data lead to immediate prestige [18, 23], but can result in the accumulation of longer-term payoffs as well [16, 24]. This comes in the form of citations, successful replications, and the remixing of data sets. A paywall is not only likely to limit the diffusion of the data set in the scientific community, but limit the number of derivative works as well.

## Methods

### Agent-based Simulation

All agent-based simulations are conducted in NetLogo 6.1.0 (Center for Connected Learning and Computer-based Modeling, Evanston, IL). This instantiation of the PD *n*-person iterative model [25] was created by Dr. Uri Wilensky [26], and is licensed under the CC-BY-NC-SA 3.0 license. All modifications to the code have been made publicly available at https://github.com/Orthogonal-Research-Lab/Models-for-Data-Reuse/tree/master/Agent-based%20Model. The world sizes for all simulation runs is 32×32 with a patch size of 8. This size was chosen so as to provide space for each agent to interact freely with all other agents, but not so large as to introduce spatial variation into the results. Each condition consists of 40 agents (20 agents per strategy). In the cooperate-defect hypothesis, 20 agents pursue a pure strategy of “cooperate”, while the other 20 pursue the pure strategy of “defect”. Strategies are evaluated in terms of average payoff, which is averaged across agents for each iteration of the model.

### Experimental Design, Hypotheses, and Payoffs

For each hypothesis, five strategy combinations each consisting of two distinct strategies are implemented: cooperation and defection (non-cooperation), cooperation and random (mixed cooperation and non-cooperation), cooperation and unforgiving (free-riding always results in subsequent non-cooperation), defection (non-cooperation) and unforgiving (free-riding always results in subsequent cooperation), and defection (non-cooperation) and random (mixed cooperation and non-cooperation).

Each hypothesis (consisting of a set of five combinations) are defined by a unique payoff matrix. This payoff matrix is structured based on the hypothesis being modeled. Three distinct conditions were used in this study. Hypothesis A (Table 1) assumes the highest payoff is for mutual cooperation, a sizable payoff for one-way free-riding, and a marginal payoff for mutual non-participation. Hypothesis B (Table 2) assumes the highest payoff is for one-way free-riding, a sizable payoff for mutual cooperation, and a marginal payoff for mutual non-participation. Hypothesis C (Table 3) assumes the highest payoff is for mutual non-participation, a sizable payoff for one-way free-riding, and a marginal payoff for mutual cooperation.

**Table 1.** Payoff matrix for Hypothesis A.

|  |  | **Transformer** |  |
| --- | --- | --- | --- |
| **Producer** |  | **Gain** | **Lose** |
|  | **Gain** | 3 | 0 |
|  | **Lose** | 2 | 1 |

**Table 2.** Payoff matrix for Hypothesis B.

|  |  | **Transformer** |  |
| --- | --- | --- | --- |
| **Producer** |  | **Gain** | **Lose** |
|  | **Gain** | 2 | 0 |
|  | **Lose** | 3 | 1 |

**Table 3.** Payoff matrix for Hypothesis C.

|  |  | **Transformer** |  |
| --- | --- | --- | --- |
| **Producer** |  | **Gain** | **Lose** |
|  | **Gain** | 1 | 0 |
|  | **Lose** | 2 | 3 |

In modeling the three hypotheses for the agent-based model, we must consider how the players (producer and transformer) interact. There are four basic interactions related to payoffs: producer gains and transformer loses; producer loses and transformer loses; producer gains and transformer gains; producer loses and transformer gains. Accumulation of social capital by both data producer and data transformer results in a mutual benefit for both parties, while mutual loss means that a loss of social capital by both data producer and data transformer. Sharing data for reuse is potentially a win-win transaction, but only if both parties recognize the payoff in social capital. As we will see in the three different hypotheses highlighted in the analysis, payoffs are highly contingent on motivations and behaviors of both the data producer and the data transformer. The greatest potential payoff occurs when these are coordinated.

### IGT Model

In IGT, one unit of payoff expressed as 1 in classical PD is calculated as

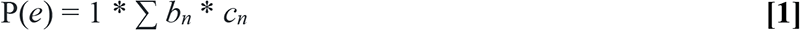

where *P(e)* is the probability of an empirical observation directly translating into a payoff, *b*_*n*_ are the collective inputs of the belief units and *c*_*n*_ are the collective inputs of the cultural constraint units. The payoff matrix is determined by the probability of an empirical observation *P(e)* being either above or below the pre-defined threshold (*e*_*T*_). *P(e)* is generally expressed as a joint probability *(e_1,2,…n_)*. An example using the joint probabilities over time of share (*e*_*1*_) and no share (*e*_*2*_) are shown in Table 4. The value of *e* for a given threshold (*x*_*T*_) is arbitrarily set at 0.5 (mean of a normal distribution), but can be any value between 0.0 and 1.0.

**Table 4.** Payoff matrix for ideological game theory.

|  | Transformer |  |  |
| --- | --- | --- | --- |
|  |  | Share | No share |
| | Share | if $e_{1,2} > 0.5$, 0 | if $e_{1,2} < 0.5$, 0 |
| | No share | if $e_{1,2} > 0.5$, 0 | if $e_{1,2} < 0.5$, 1 |

In general, the joint probability is greater when the collective influence of ideological and cultural constraint units are stronger. The general effect of ideological influence is to blur the distinction between discrete empirically-based decisions. In the above payoff matrix, the distinction between “share” and “no share” is only clear when ideological influence is above or below threshold contingent on the payoff in question.

## Results

### Data sharing as PD Game

We can model data sharing and recycling as an instance of the Prisoner’s Dilemma (PD) game. The basic payoff matrix (Table 5) can include various payoff structures for different access arrangements, but the basic set of interactions are the same. If both the producer and transformer agree to a CC-BY-NC (non-commercial) arrangement, then in principle their transactions result in a Nash equilibrium between players [27] and provides the basis for a sustainable open access community. As we will see, however, it is only one of many possible outcomes that depends on the culture in which both producer and transformer are embedded.

**Table 5.**
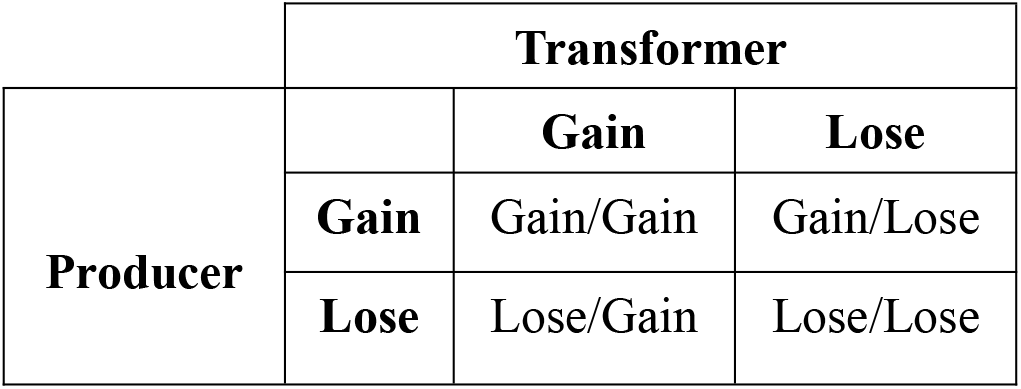
Basic structure of the payoff matrix for interactions between producer and transformer. “Gain” and “Lose” payoff categories represent the social and financial utility for the act of engaging in the open exchange of data.

|  | Transformer |  |  |
| --- | --- | --- | --- |
|  |  | Gain | Lose |
|  | Gain | Gain/Gain | Gain/Lose |
|  | Lose | Lose/Gain | Lose/Lose |

Intuitively, one might assume that the relationship between producer and transformer is one that favors the transformer only. A more cynical worldview would describe the relationship as a parasitic one, with producer being the only creator of value. In light of this, let us consider the 2×2 payoff matrix as being representative of the potential for mutual gain (Table 6). The typical application of the Prisoner’s Dilemma involves each player acting independently, but according to a common set of principles. Co-operation occurs when both the producer and transformer have been acculturated into the Open Science community.

**Table 6.**
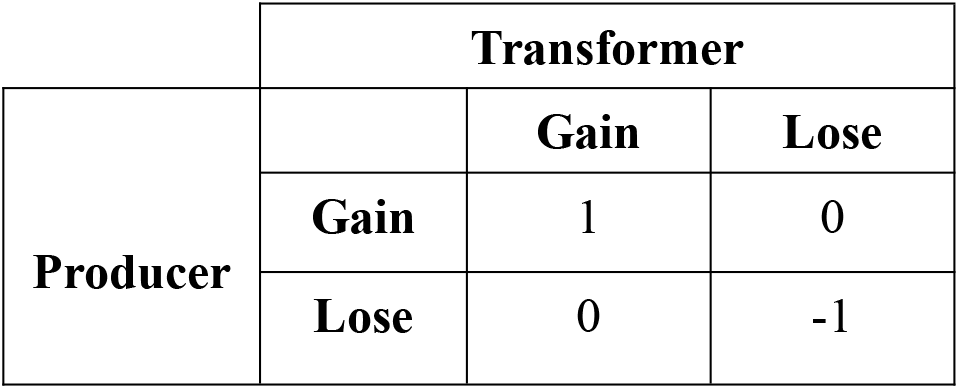
Payoff matrix if both the producer and transformer decide to share and reuse data, respectively. “Gain” and “Lose” payoff categories represent the social and financial utility for the act of engaging in the open exchange of data.

An asymmetric case in which one player loses utility for engaging in data sharing results in a payoff of zero is shown in Table 6. Figure 2 demonstrates the payoff matrix structure shown in Table 6. The most critical component of this payoff matrix involves the negative payoff when both producer and transformer lose utility by not participating in data sharing.

**Figure 2.**
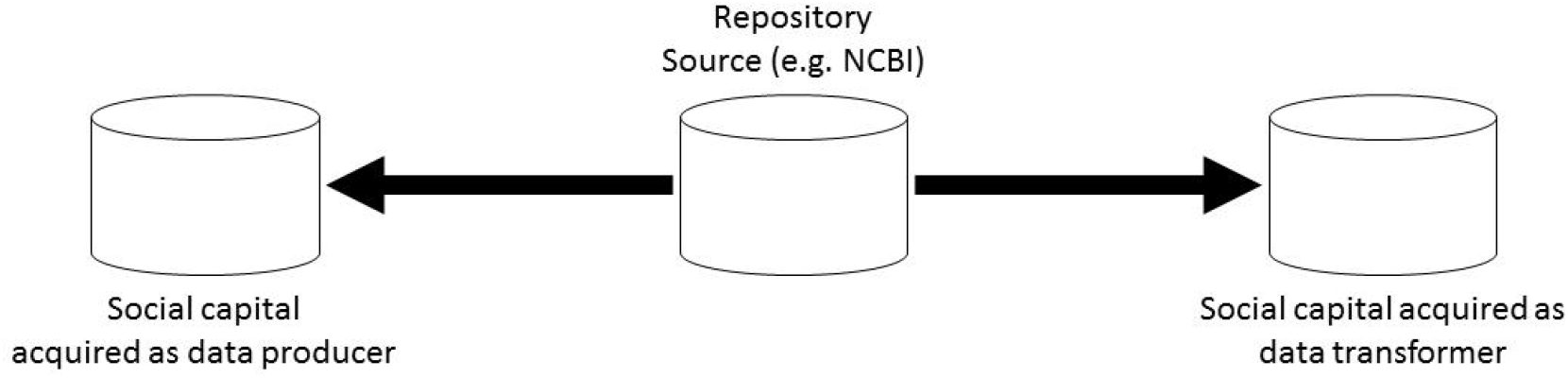
A payoff structure where data contributed to an open repository by the data producer can benefit both the primary producer and secondary transformer.

While shared principles and easy-to-use infrastructure can allow for optimal behavior, institutional skepticism about the Open Science movement can produce other outcomes. For example, let us suppose that while data sharing is considered a valuable activity, secondary use is discouraged. This is similar to the cynical view of data sharing mentioned earlier. Table 7 demonstrates the payoff matrix for this example, in which there is a negative payoff in utility whenever the producer loses, and negative to no gain whenever the transformer gains.

**Table 7.**
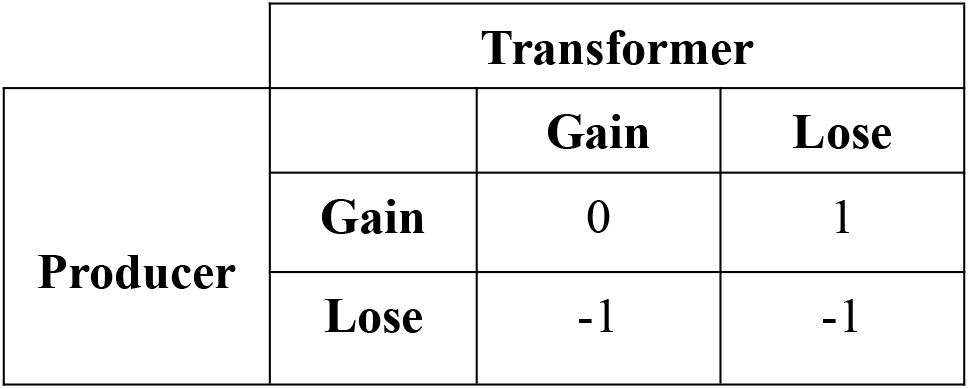
Payoff matrix if both the producer and transformer decide to share and reuse data, but gains in utility by the transformer are discouraged. “Gain” and “Lose” payoff categories represent the social and financial utility for the act of engaging in the open exchange of data.

Table 8 demonstrates what happens when sharing is discouraged entirely. As is the case in Table 7, there is no opportunity for mutual gain. In fact, the payoff matrix suggests that mutual loss is the only outcome with a payoff in utility (e.g. gain in social capital by the primary data producer). Of course, this is only true in a world where decision-makers and other people do not value Open Science. As with the other examples of PD, the utility of data sharing stems from the value assigned to both the act of sharing data and rewarding the products of data sharing. These include published datasets, meta-analysis, and the reanalysis of data.

**Table 8.**
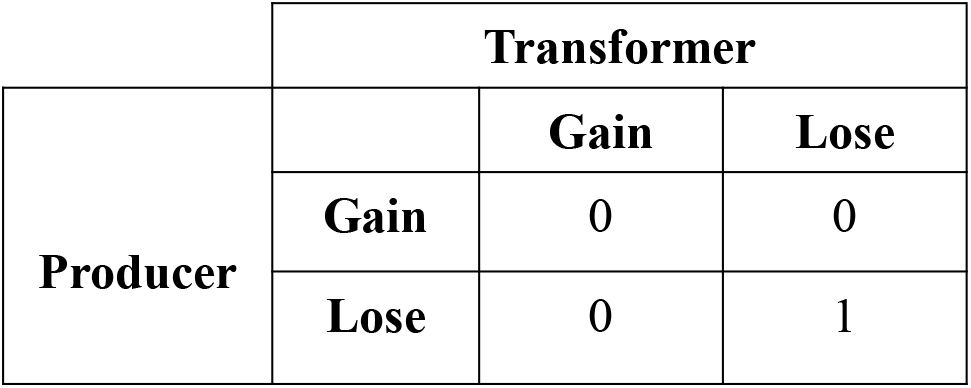
Payoff matrix if both the producer is discouraged from sharing data (both in terms of institutional and infrastructural support). “Gain” and “Lose” payoff categories represent the social and financial utility for the act of engaging in the open exchange of data.

Another feature of Tables 7 and 8 is the positive payoff for outcomes that discourage data sharing and reuse. As mentioned previously, specific payoffs are mediated by community standards, particularly when it comes to investments of time. When time is not allotted to activities such as data annotation, it can provide a motivation not to share information with the broader scientific community. In Table 7, this results in a gain for the producer (as time saved without motivation to share equals gained social capital) and a loss of utility for the transformer (as they cannot engage in their niche activity without data nor annotation information). In Table 8, this loss due to lack of proper social support results in a social capital payoff for a mutual loss of utility. In such a case, the transformer role is technically removed from the equation.

### Dynamic Simulation of Data Sharing in NetLogo

To analyze this mathematical model in the absence of an appropriate data set, a PD *n*-player iterative simulation [28] was conducted using NetLogo 6.1.0. An iterative model allows us to model the broader social outcomes resulting from a static payoff matrix by measuring the average payoff across a population of interacting agents over time. This type of model allows for the simulation of three hypotheses relevant to data sharing: one where cooperation is favored, one where defection (non-cooperation) is favored, and cases in which cooperation is favored but a significant amount of free-riding (taking advantage of other people’s work with no attribution or contribution) is tolerated. In this case, free-riding is equivalent to a lack of mutual cooperation. This might occur when the data producer does not enable the transformer or when the data transformer does not attribute (e.g. cite) the producer.

The Methods section describes each hypothesis and strategy combination tested in more detail. The results for the five strategy combinations per each of the three hypotheses are shown in Figures 3, 4, and 5. For all hypotheses, the graphs (Figure 3D, Figure 4D, and Figure 5D) that compare cooperate and unforgiving always exhibit the maximum payoffs for cooperation for both strategies. For the hypothesis that favors cooperation (Hypothesis A), there were several comparisons for agents playing cooperation against defect, a mixed strategy, and an unforgiving strategy. In the cooperation and random combination (Figure 3A), cooperation demonstrates a slightly higher average payoff after 50 iterations. A similar situation exists for the defect and random combination (Figure 3E), where defect demonstrates a slightly higher average payoff even before 50 iterations.

**Figure 3.**
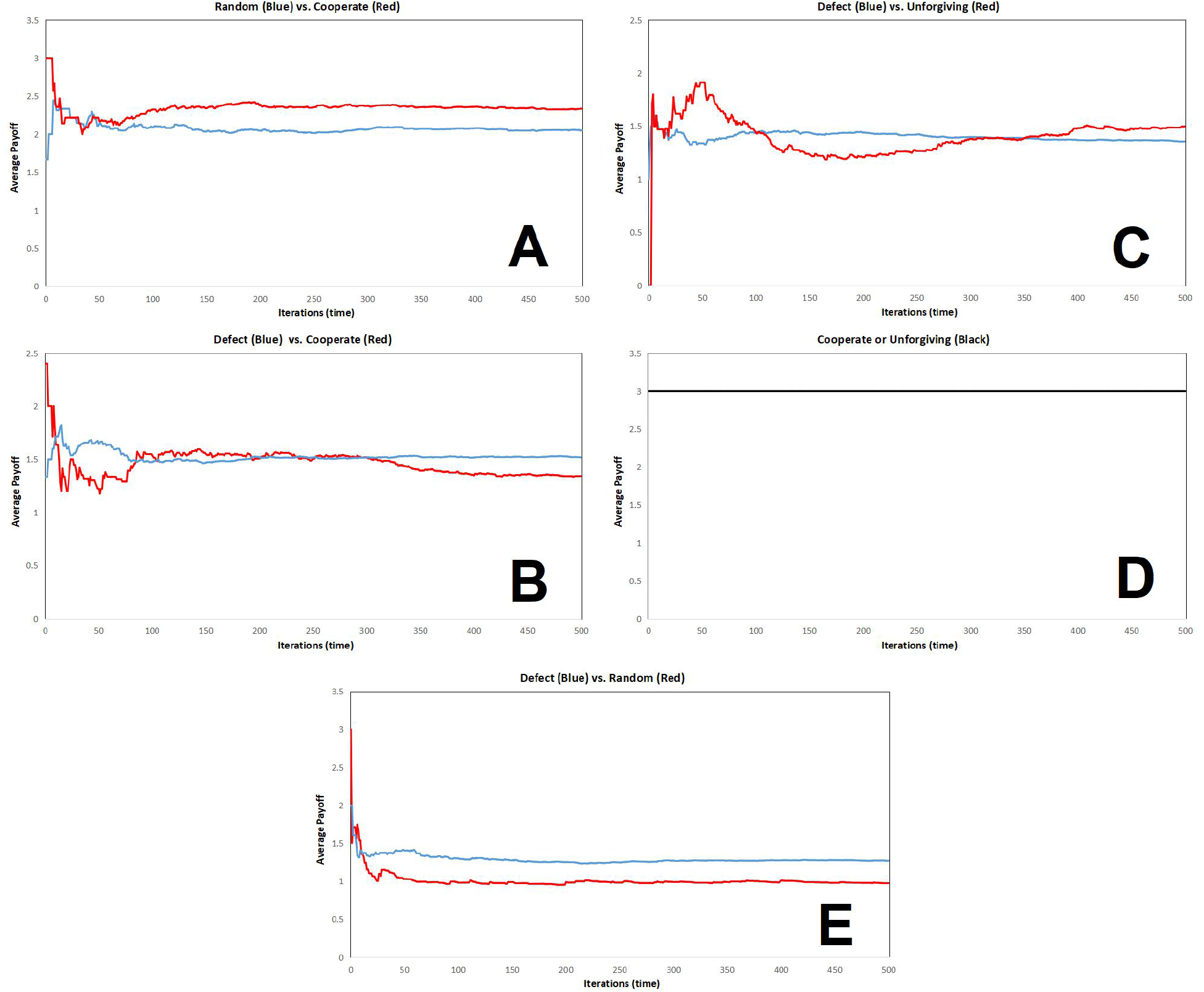
Graphs for the five combinations of head-to-head strategy comparisons for Scenario A. Panels: A, Random (blue) vs. Cooperate (red); B, Defect (blue) vs.Cooperate (red); C, Defect (blue) vs. Unforgiving (red); D, Cooperate or Unforgiving (black); E, Defect (blue) vs. Random (red).

**Figure 4.**
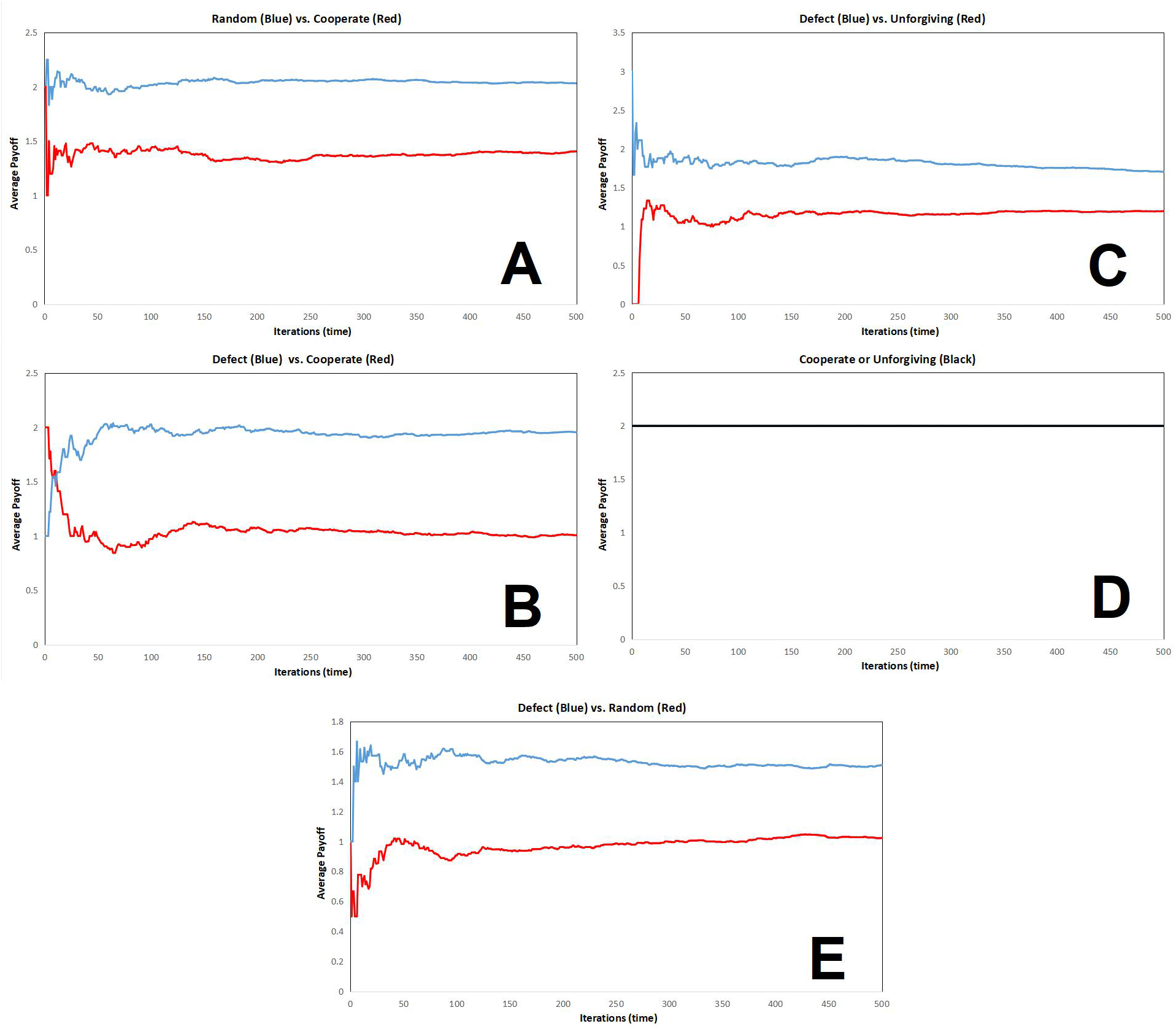
Graphs for the five combinations of head-to-head strategy comparisons for Scenario B. Panels: A, Random (blue) vs. Cooperate (red); B, Defect (blue) vs.Cooperate (red); C, Defect (blue) vs. Unforgiving (red); D, Cooperate or Unforgiving (black); E, Defect (blue) vs. Random (red).

**Figure 5.**
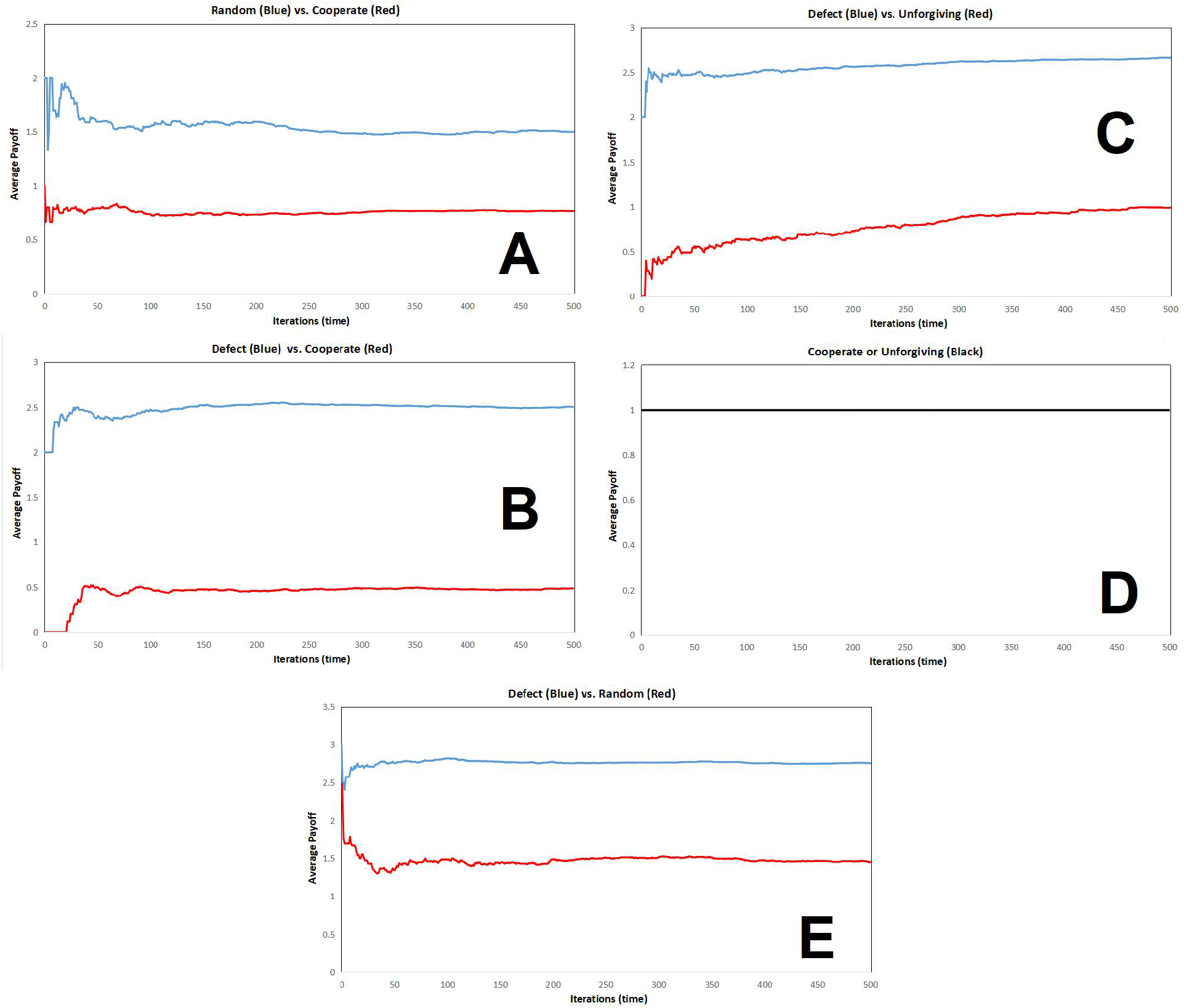
Graphs for the five combinations of head-to-head strategy comparisons for Scenario C. Panels: A, Random (blue) vs. Cooperate (red); B, Defect (blue) vs.Cooperate (red); C, Defect (blue) vs. Unforgiving (red); D, Cooperate or Unforgiving (black); E, Defect (blue) vs. Random (red).

Perhaps investing in a pure strategy (consistently share data or do not share data) provides a slightly higher payoff than inconsistently participating in data sharing. A pure defect strategy yields a higher payoff than cooperate after 300 iterations (Figure 3B). By contrast, Figure 3E shows that agents employing a pure unforgiving strategy shifts to a slightly greater payoff than a pure defect strategy after 300 iterations. Given that Hypothesis A rewards mutual cooperation, these results seem to suggest that data sharing must be both consistent over time and pervasive in order to be effective within a disciplinary community.

Now we turn to the hypothesis that favors cooperation with a significant amount of free-riding (Hypothesis B). For the first combination, the cooperate strategy always yields a lower payoff than the random strategy from the first iteration (Figure 4A). In turn, the random payoff function is far less than the defect strategy (Figure 4E). The cooperate strategy also results in a lower payoff than the defect strategy within 10-15 iterations (Figure 4B). Defect also provides a higher payoff than the unforgiving strategy (Figure 4C), while the unforgiving strategy itself yields zero payoff for the first 10-15 iterations but converges upon the defect payoff function by 500 iterations. Recall that Hypothesis B involves one-way free riding, which should be expected to yield a strong preference for non-cooperative behavior.

Lastly we examine the hypothesis that favors non-cooperation (Hypothesis C). The cooperation strategy provides less of an average payoff than the random strategy (Figure 5A). In Figure 5B, the defect strategy provides a much higher payoff than the cooperate strategy, which in this case results in an average payoff of 0 up to 20 iterations. Unforgiving is also less viable payoff-wise than the defect strategy when simulated head-to-head (Figure 5C). Finally, after a few iterations of having similar payoffs, the defect strategy’s payoff converges to a higher value than the random strategy (Figure 5E). These results tell us that when data sharing is actively discouraged, there will be a tendency to defect from potential cooperation.

### Understanding Data Sharing Using IGT

While conventional game theory provides us with a useful model of strategies employed by data sharers that approximate collective outcomes, it excludes the beliefs and cultural factors that often inhibit rational transactions. To account for these, we can use an approach called IGT [inspired by 16, 29]. IGT models account for these important exceptions by representing the empirical decisions of individuals as well as their beliefs about a particular context [30]. IGT relies upon a computational object called an *ideological perceptron* (Figure 5) that consists of three layers. The first layer features a strategy suite employed purely as a result of empirical observation (*e*_*n*_). These are the same strategies found in a conventional game theory model. The second layer models individual beliefs that might influence the empirically-based strategies (*b*_*n*_). There beliefs may provide a conditional quality to the deployment of empirically-based strategies. The third layer provides discrete cultural constraints that have a direct influence upon beliefs and an indirect influence on how empirical strategies are expressed (*c*_*n*_).

A demonstration of an IGT model as applied to the data sharing context is shown in Figure 6. In this model instance, units in all three layers can be specified with a specific semantic label, representing some human behavior or institutional factor. While it might be difficult to quantify the relative effect of each behavior/factor, this model provides at least enough information to derive payoffs and context-specific settings (such as an academic department or interdisciplinary working group). Figure 6 also shows multiple inputs converging on single units. These are dealt with in the following manner: contributions from one layer are integrated into the response of each unit in the next layer by a summation rule common to neural networks. If the summed values exceed a given threshold value (between 0 and 1), then they are active. For example, if cultural constraints act strongly enough on a given belief, that belief can be rendered irrelevant. By contrast, when there are few if any cultural constraints, new beliefs can be expressed freely.

**Figure 6.**
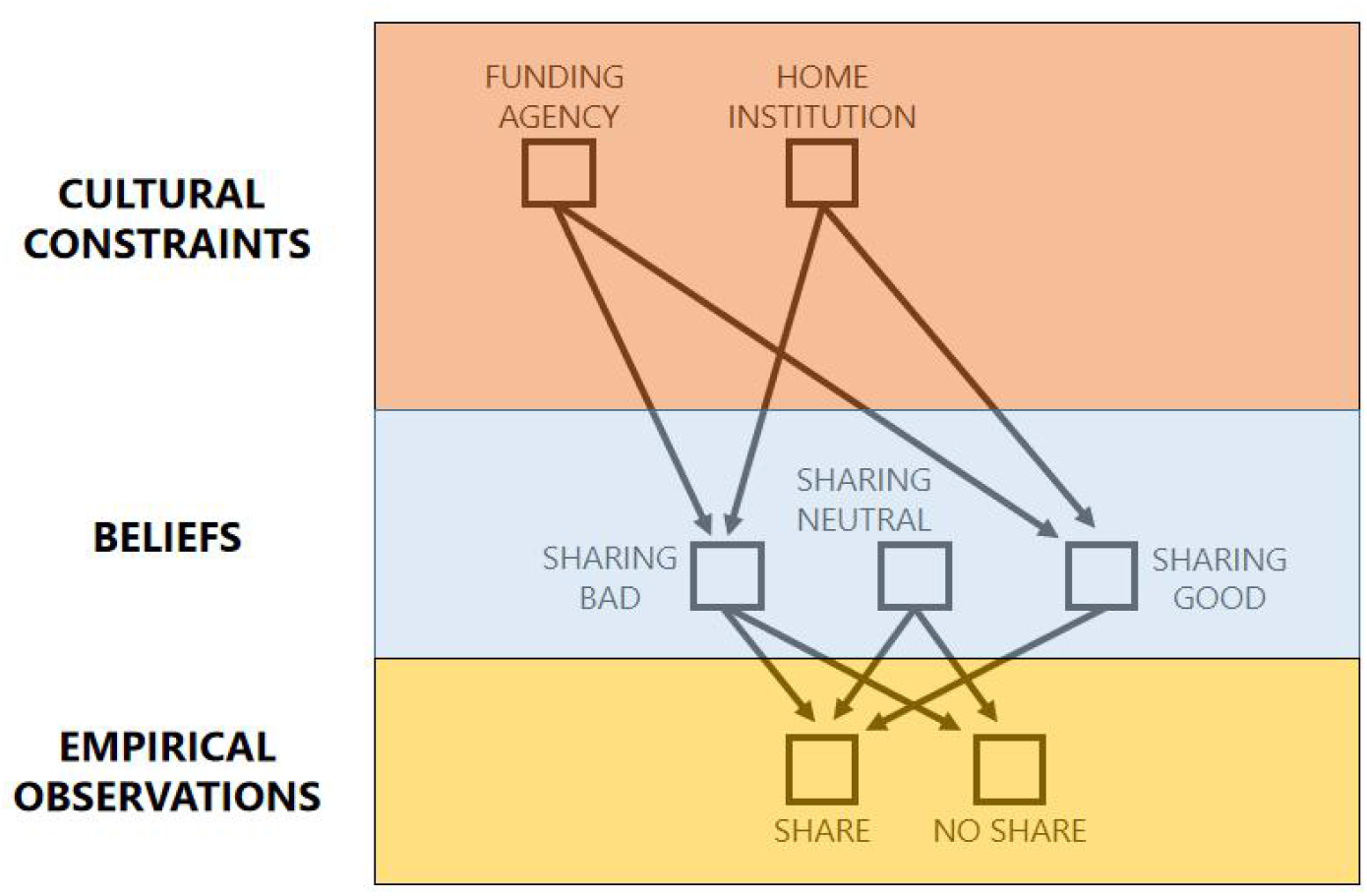
Basic configuration of the ideological perceptron, which models empirical observations, beliefs, and cultural constraints used in ideological game theory.

These new conditions allow us to reformulate our payoff matrix in a way that incorporates this new information. A main difference between classical PD and IGT is that payoffs are expressed as conditional statements. Each strategy is based on a payoff (*e*), and is represented in classical PD by a scalar value. This value is acquired by a producer or consumer through observing empirical signals and then acting upon them. In the IGT model of data sharing, empirical observations are the product of a joint distribution (*e*_*1,2*_) consisting of sharing (*e*_*1*_) and no sharing (*e*_*2*_). The distribution of sharing and no sharing is based on the degree to which beliefs (which are in turn influenced by cultural constraints) influence the empirical observation. These contributions from different cultural constraints and beliefs influence the payoff through a simple rule (see Methods). One advantage of ideological game theory is that we can represent sub-optimal behavior, or behaviors that do not conform to the most efficient outcome [31]. It also allows us to model multiple sets of community norms in the same context, which often leads to sub-optimal outcomes. This might explain the pernicious unexplained factors in the failure of communities that initially have a noble mission.

## Discussion

Given both the conventional and ideological game theory models, we can now discuss in a less formal manner which source of variation might influence decision-making and the emergence of a data sharing (or CC-BY) economy. While limited by the lack of primary data, the models presented here represent a first step toward a theoretical characterization of the social dynamics within open academic communities. The creation of such an economy is necessary for a stable and consistent culture of data sharing to emerge, particularly in institutions and disciplines where it is not already self-evident.

In general, this theoretical approach fills a gap in the literature by introducing two novel approaches (investigations of sociocultural dynamics and agent-based modeling) to the problem of understanding open data communities. The application of the classical PD model motivates the application of an agent-based model. The findings of the agent-based model confirm that as we change the incentive structure for given behaviors, we will observe expected behaviors. For example, free-riding and institutional discouragement of data sharing negatively impact the mean behavior in the community. There are, however, some interesting wrinkles when we consider the dynamics of simulated behaviors. Payoffs for certain behaviors tend to compound over time, which may not have a significant effect in a static context but shift collective behaviors in a longer-term context. This points to the importance of considering the long-term effects of incentives and enforcement of behavioral norms in a given community, and is not often apparent in the available empirical data.

Additionally, the *n*-player iterative and IGT models explore two sides of the same set of issues. While the IGT model does not provide us with concrete results with respect to data sharing, it does provide a framework for considering interpersonal and institutional biases in a way that are not possible with empirical observations nor existing models. The use of social capital as the standard currency of payoffs allows us to consider how these factors subtly influence collective behaviors in real-life data sharing communities. The consideration of social capital as a behavioral motivator suggests that financial incentives or top-down mandates may not be enough to enforce desired behaviors.

### Limitations and Conclusions

There are a number of limitations to this approach, in addition to being the inspiration for a number of empirical studies that would confirm what has been presented here. The PD *n*-player game serves as a first step towards model validation. Yet these findings also suggest that full cooperation may not be a necessary condition of building stable and sustainable communities of practice. There are two main takeaways from the agent-based model. The first is that data sharing must be both consistent over time and pervasive in order to be effective within a disciplinary community. Secondly, when data sharing is actively discouraged, there will be a tendency to defect from potential cooperation. In both cases, the agent-based model confirms that larger cultural change is necessary for a sustainable data sharing community. A third but less clear finding from these modeling experiments is that conventional economic free-riding may be tolerated at much higher levels than might be imagined by some producers of data [32]. The assumption that data production and consumption is reciprocal may also not be as straightforward as can be characterized in a simple model.

Our understanding of data sharing would benefit from a more precise definition of social capital with respect to data sharing. In particular, does social capital arise from a reciprocity transaction based on norms, or does social capital accruement arise from group membership (e.g. social solidarity)? Pigg and Crank [33] consider a number of potential mechanisms for the production and attainment of social capital in such communities. Another intriguing possibility is that existing networks formed on the basis of personal relationships or disciplinary alliances determine participation in data sharing communities and the allotment of social capital. In terms of confirming the results of the PD, *n*-player iterated PD, and IGT models, it would be good to align an empirical data set with various strategies utilized in our theoretical constructs.

Even when policy is used to encourage open data, there are still relatively few open data sets in certain fields such as medicine. Sometimes, even the cultivation of sharing-friendly cultural norms are not enough. For example, compared to older scientists younger scientists are more amenable to sharing data, but in actuality make their data less available [34]. This suggests policy is helpful but not sufficient for encouraging data sharing. Klump [20] suggests that the cultural phenomenon of gift-giving is crucial to our understanding of how participation in open data sharing communities can be encouraged. Widely observed in more traditional scientific communities [33], gift-giving might consist of exchanging information for recognition, and exemplified through citation or co-authorship. A form of gift-giving is utilized in open source software communities [35] to orient relationships in such communities towards contributions towards a public good. Gift-giving can enforce reciprocity through a formal feedback mechanism that enables future obligations [36]. Not only does the gift lead to amplifying social capital, it increases one’s ability to convert scientific raw materials into professional credibility [37].

### Towards a CC-BY Economy

This paper contributes to our understanding of how a CC-BY (Creative Commons Attribution) economy might become sustainable, particularly with respect to properly valuing and rewarding openly-available research outputs. While some CC licenses limit the ability to remix and use for commercial purposes [38], the purpose of our model is to demonstrate the role of social capital in driving interactions between research parties. A more pessimistic worldview suggests substantial cultural change needs to occur before data sharing and secondary use becomes a dominant strategy. Indeed, institutional culture and technical infrastructure can affect the outcomes predicted by this analysis. To summarize these challenges, we can look to the literature to understand the current state of academic value, norms, and economic mental models that affect decision-making among contemporary scientists.

To fully appreciate how the PD and IGT models are applicable, we must go beyond the classical economic view and establish an econosemantic one. We must first understand the economic conditions under which data is more or less likely to be shared. These conditions should then be understood through the lens of econosemantics, in this case the meaning of data sharing in a given research community. The classic view demonstrates that data sharing is constrained by a search cost function, which is currently hindered by a scattered resource landscape [39]. Resources and infrastructure that lessen the search cost often allows for the problem of data sharing to be understood using a cost-benefit framework [40]. The classic view also relies on a utilitarian view of data [41] that lends itself to the PD model.

This leads us to the econosemantic framework, which consists of two components: value construction and value translation. The first component (value construction) combines economic efficiency with community norms and cultural practice. In terms of efficiency, the literature suggests that digital tools [42], formal policy frameworks [43, 44] and public funding mandates [45] can add value to the practice of data sharing. Data sharing is encouraged the most by developing a set of shared norms in concert with software infrastructure [46]. The development and/or evolution of specific academic fields can act to constrain data sharing practices, as practices associated with a specific discipline do not often translate to interdisciplinary contexts [47]. On the other hand, the development of an interdisciplinary data sharing culture can also enable new scientific paradigms [43].

The second component of the econosemantic framework (value translation) can be identified in the open data sharing literature, and revolves around the dynamics of exchange. In open data sharing, exchange does not revolve around a single currency, but rather revolves around instances of social capital. In [48], it is suggested that a credit and incentives system can be established through various social signals and time-saving guarantees. Social incentives for sharing data can involve the currency of status-seeking through authorship [48], endorsement by an authority in a given discipline [49], or rewarding citation counts for a shared data set [50]. As part of a virtuous cycle, the practice of open data sharing itself can serve to increase the credibility of research articles and their authors [41]. Even when data is openly available, econosemantic value does not preclude its conversion into social capital that personally benefits researchers that make the appropriate investment [51].

Therefore, the greater question is what a CC-BY economy would look like, particularly in a time when the value of research outputs are in flux, is still to be determined. While game-theoretic models such as PD lead to stable cooperation, iterated PD models can also result in extortion behaviors [52]. The cornerstone of a CC-BY economy rests upon the convertibility of social capital and intangible benefits of communal maintenance to some form of financial reward. A related factor is explicitly encouraging investments in annotating and the analysis of data. Another critical component of a CC-BY economy is greater recognition of the symbolic and semantic precursors of value. This ranges from recognizing the benefits of open access on the diffusion of research to changing the language of science to reflect desirable but undervalued contributions. Typically, components of the traditional academic economy come imbued with values that discourage open data, and thus the existing economic rewards and social practices within this economy have an inherent bias against Open Science practices [53, 54]. All too often, this leads us to the scenario in Tables 3 and 4 than the one in Table 2. It is up to us to shape the future, and the principles of Open Access enable that way forward.

## Notes

#### Summary of Updates

The title and abstract have been modified to reflect content changes. An agent-based model was also added to the paper, with changes to the Introduction, Figures, Discussion, and References. Three figures have been added to demonstrate the results of the agent-based simulation. A Methods section has also been added to reflect the additional experimental work.

